# Harnessing Deep Learning to Analyze Cryptic Morphological Variability of *Marchantia polymorpha*

**DOI:** 10.1101/2023.05.31.543177

**Authors:** Yoko Tomizawa, Naoki Minamino, Eita Shimokawa, Shogo Kawamura, Aino Komatsu, Takuma Hiwatashi, Ryuichi Nishihama, Takashi Ueda, Takayuki Kohchi, Yohei Kondo

## Abstract

Characterizing phenotypes is a fundamental aspect of biological sciences, although it can be challenging due to various factors. For instance, the liverwort (*Marchantia polymorpha*), a model system for plant biology, exhibits morphological variability, making it difficult to identify and quantify distinct phenotypic features using objective measures. To address this issue, we utilized a deep learning-based image classifier that can handle plant images directly without manual extraction of phenotypic features, and analyzed bright-field images of *M. polymorpha*. This dioicous plant species exhibits morphological differences between male and female wild accessions at an early stage of gemmaling growth, although it remains elusive whether the differences are attributable to sexual dimorphism or autosomal genetic variation. To dissect the genomic factors, we established a male and female set of recombinant inbred lines (RILs) from a set of male and female wild accessions. We then trained deep-learning models to classify the sexes of the RILs and the wild accessions. Our results showed that the trained classifiers accurately classified male and female gemmalings of wild accessions in the first week of growth, confirming the intuition of plant researchers in a reproducible and objective manner. In contrast, the RILs were less distinguishable, indicating that the differences between the parental wild accessions arose from autosomal variations instead of sexual dimorphism. Furthermore, we validated our trained models by an “explainable AI” technique that highlights image regions relevant to the classification. Our findings demonstrate that the classifier-based approach provides a powerful tool for analyzing plant species that lack standardized phenotyping metrics.

## Introduction

Plant phenotypes are influenced by genetic and environmental factors, resulting in differences in shape, color, and generation of specific organs. Although the characterization of such phenotypes is a fundamental aspect of plant biology, it often poses considerable challenges. For instance, the liverwort (*Marchantia polymorpha*), an emerging model system in plant biology (Bowman et al. 2022; Kohchi et al. 2021), has phenotypic traits that are not easily quantifiable. *M. polymorpha* grows as a flattened creeping thallus that periodically bifurcates from the apical notch (Shimamura 2015; Solly et al. 2017). The continuous architecture is variable and thus hinders the identification and quantification of distinct phenotypic features using objective measures. This difficulty makes potentially significant phenotypes cryptic, which means they are recognizable to the human eye but not available for quantitative and statistical analysis. Therefore, developing a method to handle such cryptic phenotype features would facilitate the understanding of *M. polymorpha* and other plant species that differ from well-characterized model organisms.

Phenotypic characterization and comparison using handpicked features, such as the aspect ratio of leaves, is a classic yet effective method. However, such features often lack the necessary expressiveness to describe diverse biological forms. This has led to the development of sophisticated shape and color analysis techniques, often utilizing computer-assisted image analysis (Chitwood and Sinha 2016). For example, Fourier-based analysis allows the decomposition of an arbitrary contour shape into a sum of periodic components with different frequencies, and has been applied to characterize cell and tissue shapes. Additionally, the characterization of surface properties such as color and texture has been an active research field to understand, for example, floral evolution driven by the color preferences of insects (Ohashi et al. 2015). However, it remains challenging to analyze shape, color, and texture information in a unified manner. Consequently, the discriminative ability of phenotyping methods for image datasets is often inferior to that of the human eye. A research direction for resolving this issue would be to employ neural-network-based image classifiers, which have produced promising results (Singh et al. 2018). Furthermore, Akagi and his colleagues adopted not only deep learning-based classifiers but also “eXplainable AI (XAI)” techniques to visualize the reasons behind diagnosis of calyx-end cracking in persimmon fruits (Akagi et al. 2020). This demonstrates the potential of such techniques to augment cognitive capacity of researchers.

The image-classifier-based approach offers an advantage by eliminating the need for defining specific features. Deep learning models can learn complex features specialized for classification tasks, and thus they are capable of detecting cryptic morphological variability that may not be easily detectable using traditional morphological or visual assessments. However, it should also be noted that there is no guarantee that the deep learning models will use biologically meaningful features in images (Khorram et al. 2021). For example, differences in photographic condition and photographer can lead to subtle changes in color and brightness of images. If a classifier utilizes such irrelevant image features, it would not have any biological significance, regardless of the accuracy of the classification.

To maximize the above benefit and reduce the risk at using deep learning models, we devised an analysis protocol that includes a validation step of the trained models by human-interpretable feature ablations and XAI techniques to distinguish biologically meaningful classifiers from meaningless ones. Here, we demonstrated the effectiveness of our approach by resolving an open question regarding the morphological variability of *M. polymorpha*. This dioicous plant species often exhibits morphological differences between male and female plants even in accessions collected from the same site, such as Takaragaike accessions (male Tak-1 and female Tak-2) (Bowman et al. 2017) and Australian/Melbourne (Aus) accessions (Flores-Sandoval et al. 2015).

However, it remains elusive whether the differences in these wild accessions arise from sexual dimorphism or autosomal genetic variations. In this study, to dissect the genomic factors, we established male and female recombinant inbred lines (RILs) from Tak-1 and Tak-2. We then utilized deep-learning models to classify the sexes from bright-field images of the RILs and the wild accessions in gemmalings during the first week of development. To achieve this, we adopted ResNet50 (He et al. 2016), a model architecture that has been proven to be highly effective in image classification. Furthermore, we validated the trained models by an XAI technique that highlights image regions relevant to the classification. Our study paves the way for application of deep learning models for analyzing plant species that lack standardized phenotyping metrics.

## Results

### Establishment of recombinant inbred lines Rit-1/Rit-2

Common laboratory accessions, such as Tak-1/Tak-2 and Aus male and female lines, were collected as independent plants and therefore possess polymorphisms between the counterparts. To reduce autosomal polymorphisms, we generated RILs. Briefly, we first crossed male Tak-1 and female Tak-2 to obtain the F1 generation. We then inbred the siblings four times to obtain F5 generation siblings, which were named ‘Recombinant Inbred line derived from Takaragaike (Rit)’, Rit-1 (male) and Rit-2 (female) (Fig. 1A). We sequenced the whole genome of Rit-1 and Rit-2 and mapped them, together with Tak-2, to the genome of Tak-1 to call variants (Fig. 1B). Rit-1 and Rit-2 shared the same polymorphism patterns in the autosomes. 57.0% of the autosomal regions did not contain a significant number of polymorphisms between Tak-1 and Tak-2 (Supplementary Table S1), which may have resulted from a natural cross(es) before the collection of Tak-1 and Tak-2. 30.7% of the autosomal regions contained a significant number of polymorphisms between Tak-1 and Tak-2 as well as that between Tak-1 and Rit-1 or Rit-2 (Supplementary Table S1), indicating that this region was derived from Tak-2 and the rest (12.3%) from Tak-1. These data indicate the establishment of Tak-1/Tak-2-derived RILs.

**Fig. 1.**
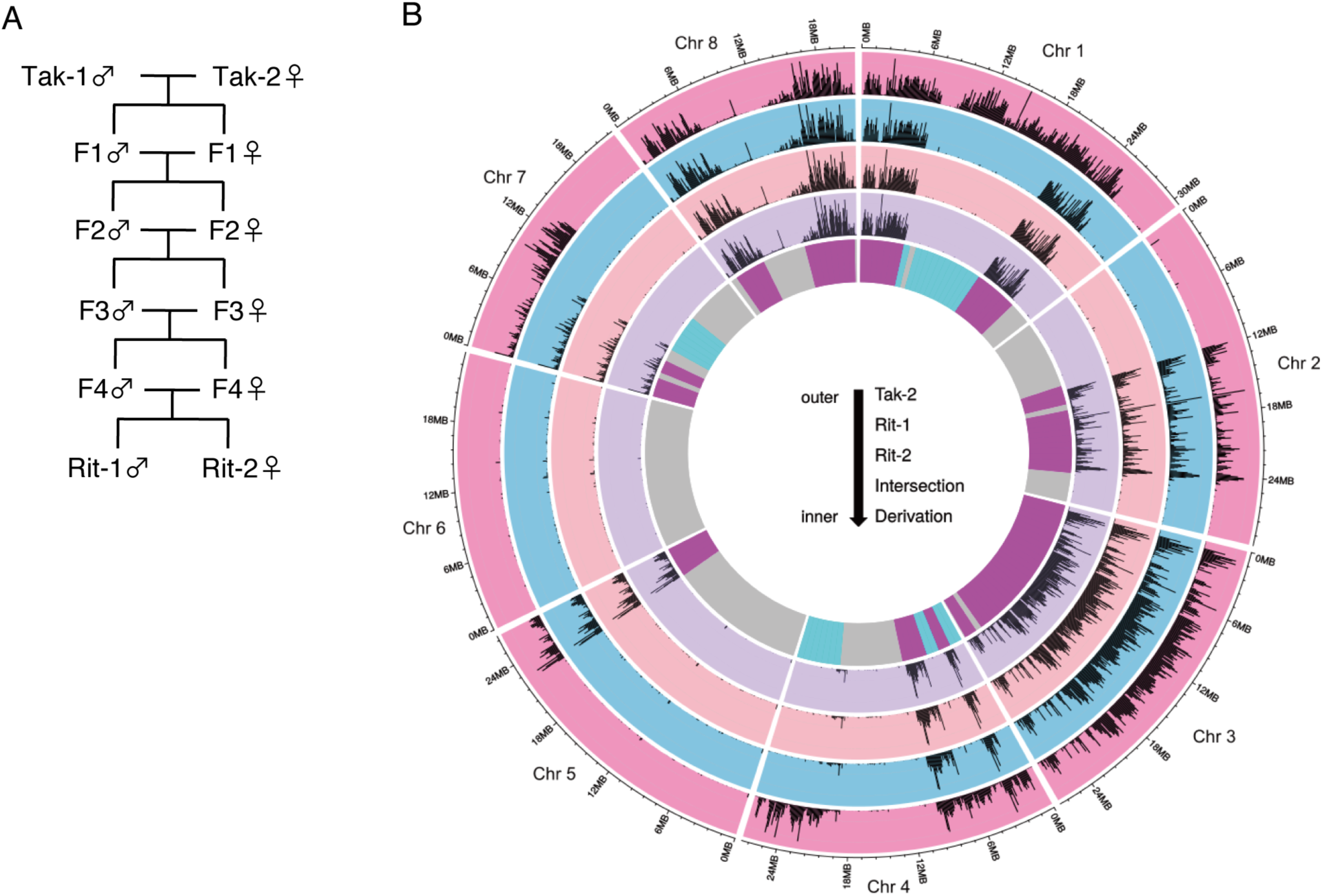
Establishment of recombinant inbred lines. (A) Inbreeding scheme to generate Rit-1 (male) and Rit-2 (female). (B) Circos plot showing the derivation of genomic regions in RILs’ autosomes. Outer tracks (tracks 1-4) show polymorphism frequencies compared to Tak-1: track 1, Tak-2; track 2, Rit-1; track 3, Rit-2; track 4, intersection of Tak-2, Rit-1 and Rit-2. The most inner track depicts predicted derivation of genomic regions in Rit-1/Rit-2. Cyan, regions derived from Tak-1; magenta, those from Tak-2; gray, those shared between Tak-1 and Tak-2.

### Deep learning for sex classification of Marchantia polymorpha based on gemmaling images

To acquire images of *M. polymorpha* gemmalings, we used male Rit-1 and female Rit-2, their parental male Tak-1 and female Tak-2, and male/female of another wild accession Aus. Fig. 2A shows representative images of the lines and developmental days. The number of male/female images was approximately 100/100 for the Tak-1/Tak-2 and Aus, and approximately 300/300 for Rit-1/Rit-2 (See Supplementary Table S2 for details), respectively. Images were acquired on days 0, 1, 2, 3, 4, and 7 after planting gemmae. These images were processed and used to train deep-learning image classifiers and for post-hoc analyses.

**Fig. 2.**
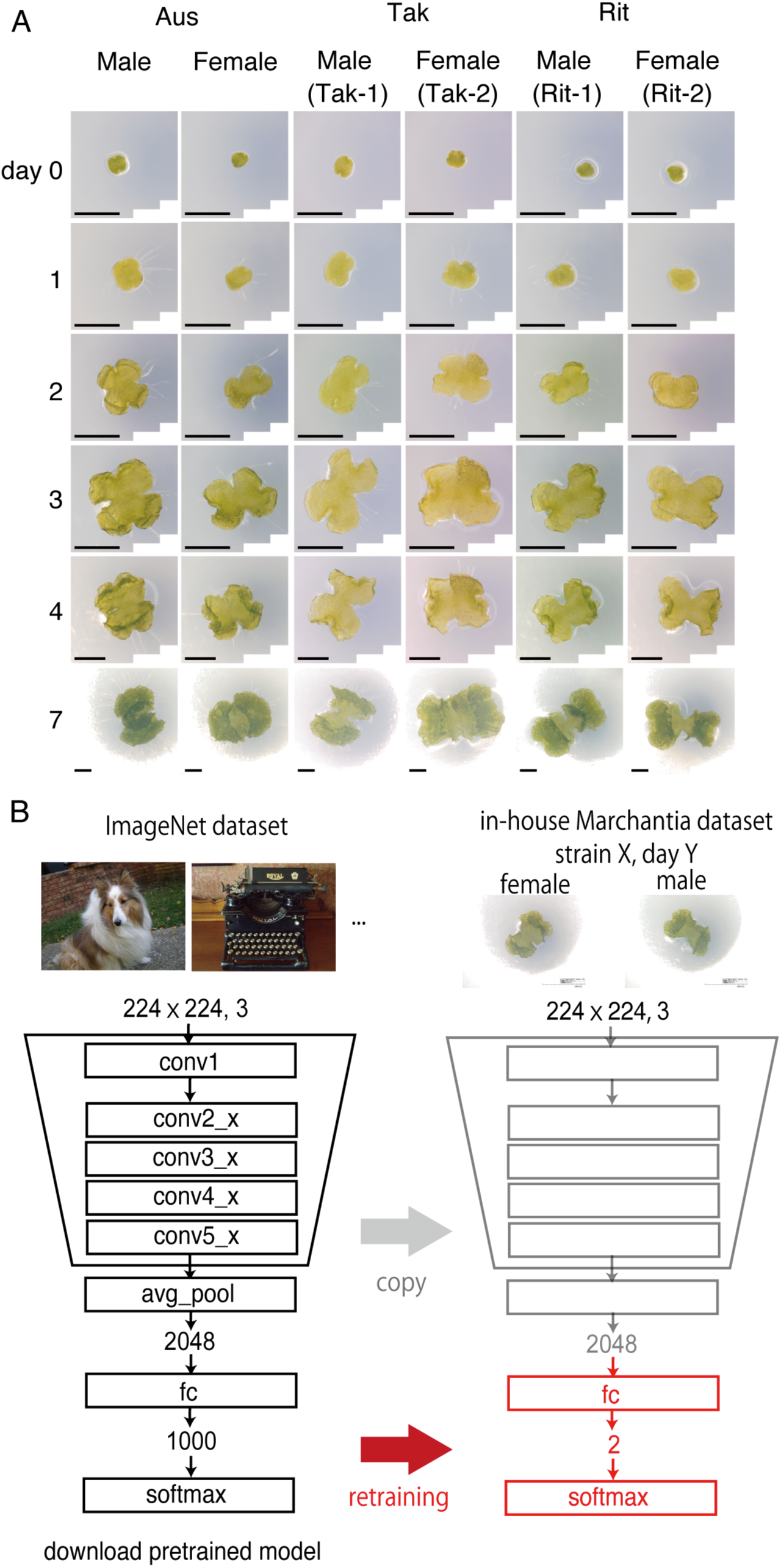
Experimental and analytical workflow. (A) Representative images of early gemmalings from Aus, Tak-1/Tak-2, and Rit-1/Rit-2. Rows: developmental day of gemmalings. Columns: accession and sex. Scale bars, 1 mm. (B) Training scheme of pretrained ResNet50 model. Only the last fully-connected layer was modified and re-trained for sex classification of gemmaling images.

Throughout this study, we utilized ResNet50 (He et al. 2016), which is a well-tested deep-learning model for image classification. The model was fed with input information, i.e., images of *M. polymorpha* gemmalings and sex labels associated with each image, to predict the sex of the new images with hidden sex labels. Although we took hundreds of images of gemmalings, the dataset size was much smaller than that of common training datasets, which typically contain millions of images. Because deep learning models have numerous parameters (e.g. 25.6 million in ResNet50), training a model with a small dataset may lead to overfitting, a situation in which a model simply memorizes all training images and corresponding sex labels instead of extracting meaningful morphological traits. To avoid overfitting, we employed a common approach for transfer learning. As illustrated in Fig. 2B, we utilized a pre-trained ResNet50 model with the ImageNet dataset (Russakovsky et al. 2015) as a starting point, and then re-trained only the last layer of the model to classify the sex of the plant (See Materials and Methods for further details about the training process) (Donahue et al. 2014; Sharif Razavian et al. 2014).

First, we quantified the simplest morphological feature, the area of the gemmalings (Fig. 3A). In Tak-1/Tak-2, Tak-2 (female) was larger than Tak-1 (male) after day 1. Aus males and females were similar in size up to day 4, but females became larger at day 7. Rit-1/Rit-2 did not differ in size throughout the examined period. The area differences between males and females were quantified by a statistical measure, unbiased Cohen’s d (Gurevitch and Hedges 2001) (Supplementary Table S3).

**Fig. 3.**
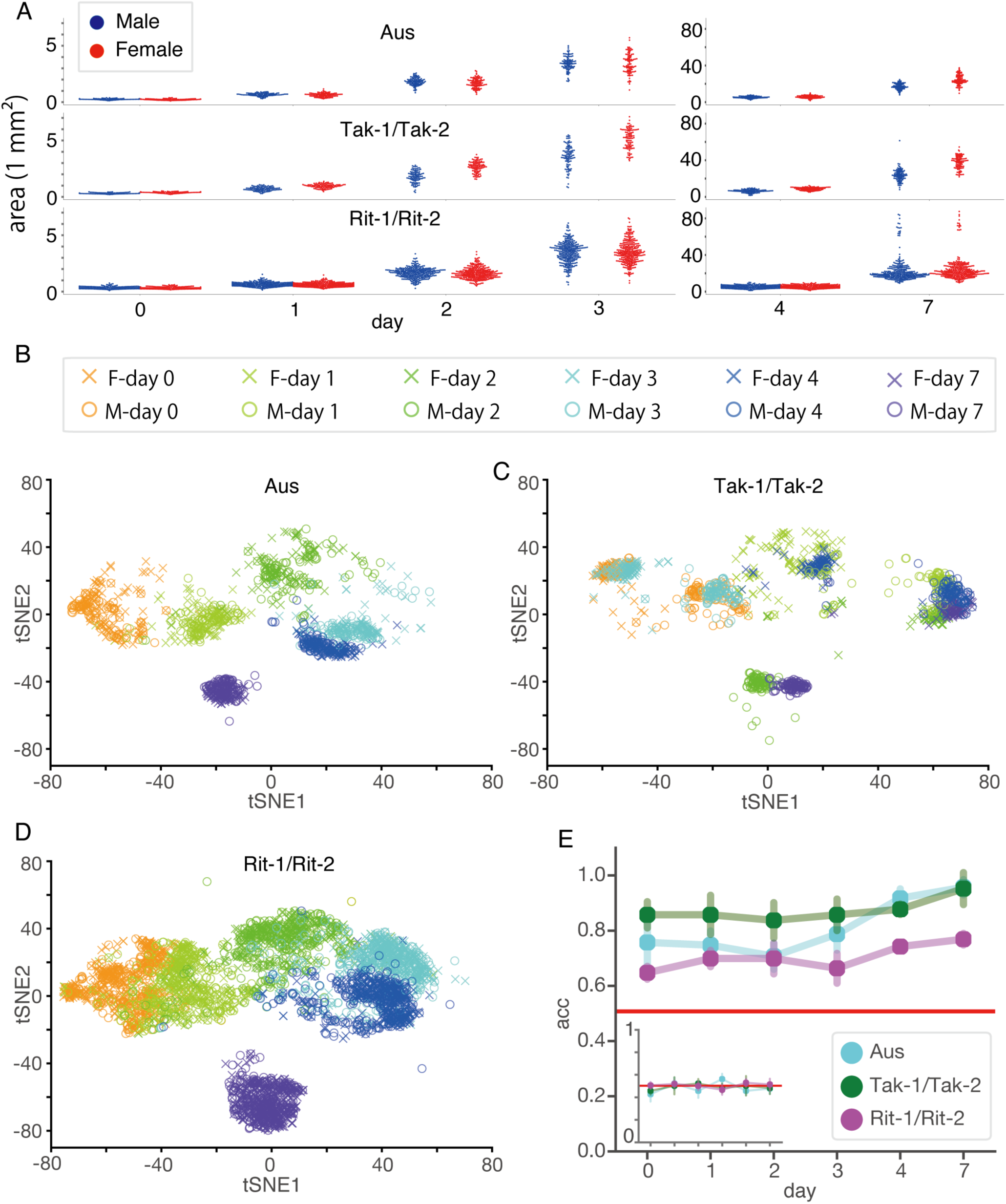
Sex classification of *Marchantia polymorpha* using the ResNet50 architecture. (A) Quantification of area of aerial part of gemmalings of Aus, Tak-1/Tak-2, and Rit-1/Rit-2. Horizontal axis represents the day after planting gemmae. Each dot represents an individual image. (B-D) Image clustering using t-SNE for Aus (B), Tak-1/Tak-2 (C), and Rit-1/Rit-2 (D). We extracted 2048-dimensional features from individual images using ResNet50 and then applied t-SNE to the feature vectors. Each point represents a different image. (E) The test accuracy of sex classification was plotted against developmental days of gemmalings. Points and bars indicate mean and SD for 5 independent trials with random training/validation/test splitting for each day. Statistical analyses were performed on day 7. Asterisks indicate a significant difference in a two-tailed Welch’s t test with Bonferroni adjustment, p<0.01; ns, not significant; n=5. The inset shows negative controls where the sex labels of the images were randomly permuted.

Nextly, we visualized the dataset by applying a nonlinear principal component analysis (PCA) technique called t-distributed stochastic neighbor embedding (t-SNE) (Maaten and Hinton 2008). Fig. 3B-D displays the clustering of all images, encompassing three lines and six developmental days. The entire dataset was clustered, but for better visualization, the results are presented in separate figures. In Aus (Fig. 3B) and Rit-1/Rit-2 (Fig. 3D), the images were grouped into clusters according to the day of development; however, those of different sexes were not distinctly placed into separate clusters. In contrast, Tak-1/Tak-2 (Fig. 3C) were separated by sex, and the images were periodically embedded according to developmental days: the first cycle (days 0, 1, and 2) and the second cycle (days 3, 4, and 7). These clustering results suggest that the morphological differences between sexes are difficult to discriminate in Aus and Rit-1/Rit-2, but are clearer in Tak-1/Tak-2.

Fig. 3E shows the classification accuracy for the test images, i.e., the accuracy for images not used in the training phase. Test accuracy, unlike training accuracy, helps to detect invalid models that simply memorize training images without learning meaningful features. For each developmental day and accession, the images were independently analyzed using different models; hence, each model addressed the binary classification of male/female. For each classification task, we conducted five independent model training sessions using randomly selected training, validation, and test images from the dataset. We used 64% of the images for training, 16% for validation, and 20% for testing. As a negative control, we trained the same model based on randomized sex labels (Fig. 3E, inset). This confirmed that the test accuracies in the negative control setting were indistinguishable from chance.

As shown in Fig. 3E, the gemmaling images of Tak-1/Tak-2, compared to those of Aus and Rit-1/Rit-2, were most easily classified, with an accuracy of over 90% on day 7. This is consistent with the t-SNE clustering (Figs. 3B-C) which suggests that Tak-1 and Tak-2 are more sexually distinguishable than the other accessions. The classification accuracy for Aus was relatively low on day 0, whereas the accuracy increased substantially after day 2, becoming comparable to that of Tak-1/Tak-2 after day 4. Meanwhile, the trained models encountered difficulty in accurately classifying the test images of Rit-1/Rit-2 and demonstrated the lowest accuracies (from 64.0±2.3% at day 0 to 76.2±2.1% at day 7) among all examined accessions, despite having a larger number of training images compared to the wild accessions. This finding objectively confirms the intuition of researchers that morphological differences between sexes in wild accessions are more distinct and these differences are diminished in the inbred lines.

### Dissecting image regions relevant to decision-making of trained deep-learning models

Deep learning models are capable of utilizing any type of visual property, such as texture, color, shape, and the presence of particular objects. However, a model may undesirably recognize irrelevant features such as background medium color, rather than sex-associated morphological differences. To investigate the features that the trained model utilized for classification, we removed certain features from the original images and newly trained the ResNet50 model with the feature-ablated images.

We conducted three types of feature ablations: (1) masking the background with black, (2) binarization into white foreground and black background, and (3) binarization and severe blurring (Fig.4A). For Ablation (1), we eliminated information from the background by filling all the background areas in black. If the model recognized features from the background, then the test accuracies decreased compared to those from the original setup. Thus, the decrease in accuracy quantifies the information from the background. For Ablation (2), we binarized images to erase the information of color and texture of the plant aerial part as well as the information in the background. For Ablation (3), we severely blurred the binarized images to render detailed contours unrecognizable.

Fig. 4B-D shows the test accuracies of the feature ablation experiments for Aus (Fig. 4B), Tak-1/Tak-2 (Fig. 4C), and Rit-1/Rit-2 (Fig. 4D). The test accuracy was reduced under most conditions after Ablation (1), suggesting that the original models utilized background information in their decision-making. Additionally, Ablation (2), which involved binary masking, resulted in further degradation of the test accuracy. This indicates that the color and/or texture of gemmalings are at least equally informative as compared to the background for classification. On the other hand, Ablation (3) did not lead to accuracy reduction, indicating that the blurred masks and the binary masks convey the same amount of information for classification. It is not surprising that the images of wild accessions were still classifiable under the most severe ablation because the gemmalings of Aus and Tak-1/Tak-2 exhibited area differences between the sexes. When comparing the accessions, the classification accuracies for Rit-1/Rit-2 were lower than those for the wild accessions and dropped to near-chance rates under Ablations (2) and (3). The results confirmed that Rit-1/Rit-2 has less sexual difference than the wild accessions.

**Fig. 4.**
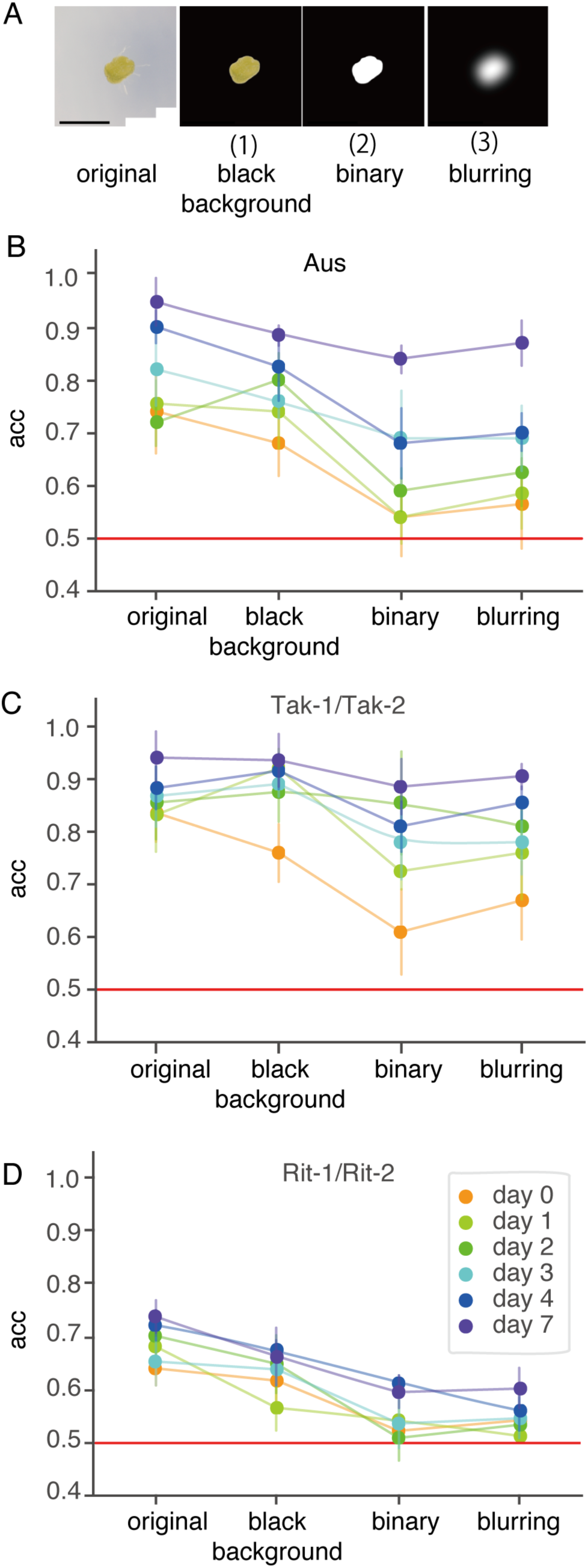
Validation of the trained classifier with feature ablation. (A) Examples of feature-ablated images. (B) Plot showing test accuracy against feature-ablation types for Aus, Tak-1/Tak-2, Rit-1/Rit-2. The same ablation procedure was applied to training, validation, and test datasets in each model-training session. The data points are grouped by developmental days, indicated by the colors. Points and bars represent mean and SD, respectively, for 5 independent trials with random training/validation/test splitting.

For further validation, it would be useful to analyze the parameters of the trained models. However, such an analysis is intractable because of the sheer complexity of deep learning models. To make these models transparent and interpretable, post-hoc model-analysis methods known as “eXplainable AI (XAI)” have been developed in the past decade. To visualize the relevant regions for the decision-making of the trained models, we employed Grad-CAM (Selvaraju et al. 2017), which is one of the most well-tested XAI methods for image classifier models. Grad-CAM generates a “class-discriminative localization map”, which is a heatmap superimposed on an input image that highlights discriminative regions containing critically important features for classification.

Grad-CAM heatmaps visualized whether sex-associated discriminative features were located in the gemmalings themselves or in the background. If the models appropriately pay attention to plant area for both sexes, class-discriminative features should appear in the gemmaling area. In contrast, the Grad-CAM heatmaps on the background instead of the gemmalings indicate that the models utilized irrelevant features.

Fig. 5 displays the Grad-CAM results. Since Grad-CAM heatmaps are computed for each individual input image, we calculated Intersection over Union (IoU) to summarize image-wise explanations across the dataset. The IoU score is a metric for measuring the degree of overlap between two regions of interest, where 1 (maximum) indicates a pixel-perfect match and 0 (minimum) indicates no overlapping area. Fig. 5A shows the IoU scores between the Grad-CAM heatmaps and the aerial parts of the gemmalings for all test images. Note that although Grad-CAM heatmaps were generated for both sexes per image, only the IoU scores for predicted sex labels are shown to focus on the decision-making of the models (See Fig. S1 for the Grad-CAM IoU scores with the medium region and the whole image area). From days 3 to 7, the Grad-CAM heatmaps overlapped with the gemmalings in both male and female images. Meanwhile, until day 1, one sex had heatmaps in the gemmaling regions whereas the other sex did not. Zero IoU values suggest that the model responded to nuisance features such as background medium color. Interestingly, all accessions exhibited a transition in the IoU distribution around day 2, although the time courses of classification accuracy were largely different among the accessions.

**Fig. 5.**
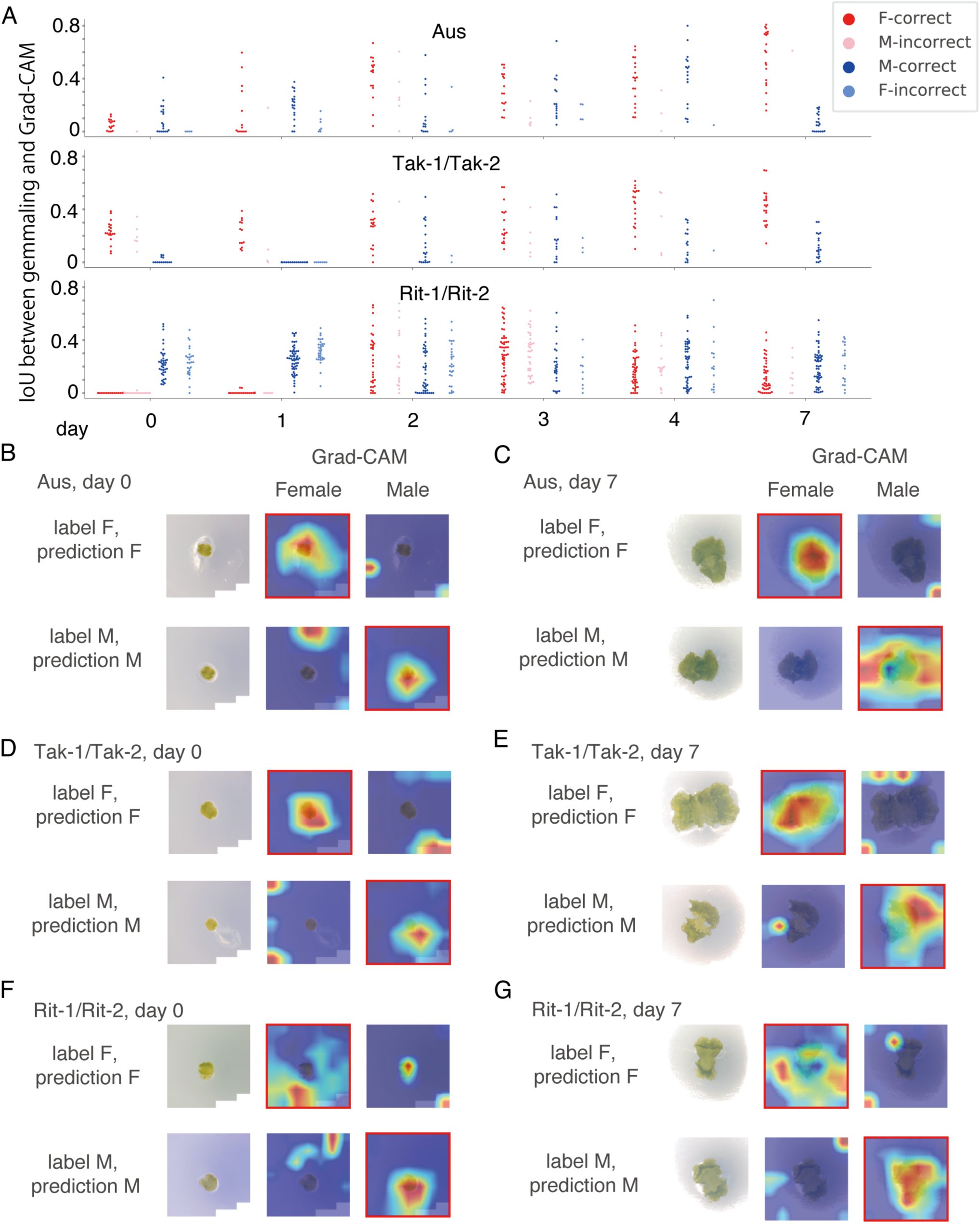
Grad-CAM heatmaps visualize the image regions relevant to classification. (A) Quantification of overlap between Grad-CAM heatmap and gemmaling with IoU score. The Grad-CAM heatmaps were computed for the predicted sex labels. Each dot represents an image. Red and blue represent correctly classified images of females and males, respectively. Pink and pale blue represent misclassified images where true labels are male and female, respectively. For each accession and day, the IoU scores for all test images are shown. (B-G) Representative Grad-CAM heatmaps for Aus day 0 (B), Aus day 7 (C), Tak-1/Tak-2 day 0 (D), Tak-1/Tak-2 day7 (E), Rit-1/Rit-2 day 0 (F), and Rit-1/Rit-2 day 7 (G).

Fig. 5B-G shows representative images of the Grad-CAM heatmaps on days 0 and 7. To avoid displaying cherry-picked results upon selecting a single representative heatmap among the 40 (Aus and Tak-1/Tak-2) and 120 (Rit-1/Rit-2) candidate images for each condition, we quantified the degree of confidence that the models had in each classification and used the images with the highest confidence scores. In line with our observations in Fig. 5A, the classifiers at day 0 often focused on the background (top center in Fig. 5F) or scratches on the culture medium (bottom right in Fig. 5D), rather than on gemmalings. On day 7, the Grad-CAM heatmaps overlapped the gemmalings in female images while rhizoids in male images of the wild accessions (Fig. 5C and E). This tendency in the wild accessions was reversed for the representative images of Rit-1/Rit-2 (Fig. 5G), although the distribution of IoU scores on day 7 did not support the reversed tendency at a dataset scale (Fig 5A).

### Tak-1/Tak-2 classifiers failed to distinguish between Rit-1/Rit-2 images

It would be interesting to examine whether the inbred lines, Rit-1 and Rit-2, resemble the parental lines Tak-1 or Tak-2. Thus, we adopted the models trained with Tak-1 and Tak-2 images to classify sex from the Rit-1 and Rit-2 images (Fig. 6). On days 0, 1, and 4, Rit-1/Rit-2 images were classified equally often as Tak-1 or Tak-2, irrespective of the true sex (Fig. 6A, B, and E). On days 2, 3, and 7, most of the Rit-1/Rit-2 images were classified as Tak-1, which is male (Fig. 6C, D, and F). That is, on all developmental days, the classification accuracy was poor. The failure of the classifiers was confirmed by Matthew’s correlation coefficient (MCC). These results indicated that the morphological differences between male and female in parental Tak-1/Tak-2 disappeared in Rit-1/Rit-2.

**Fig. 6.**
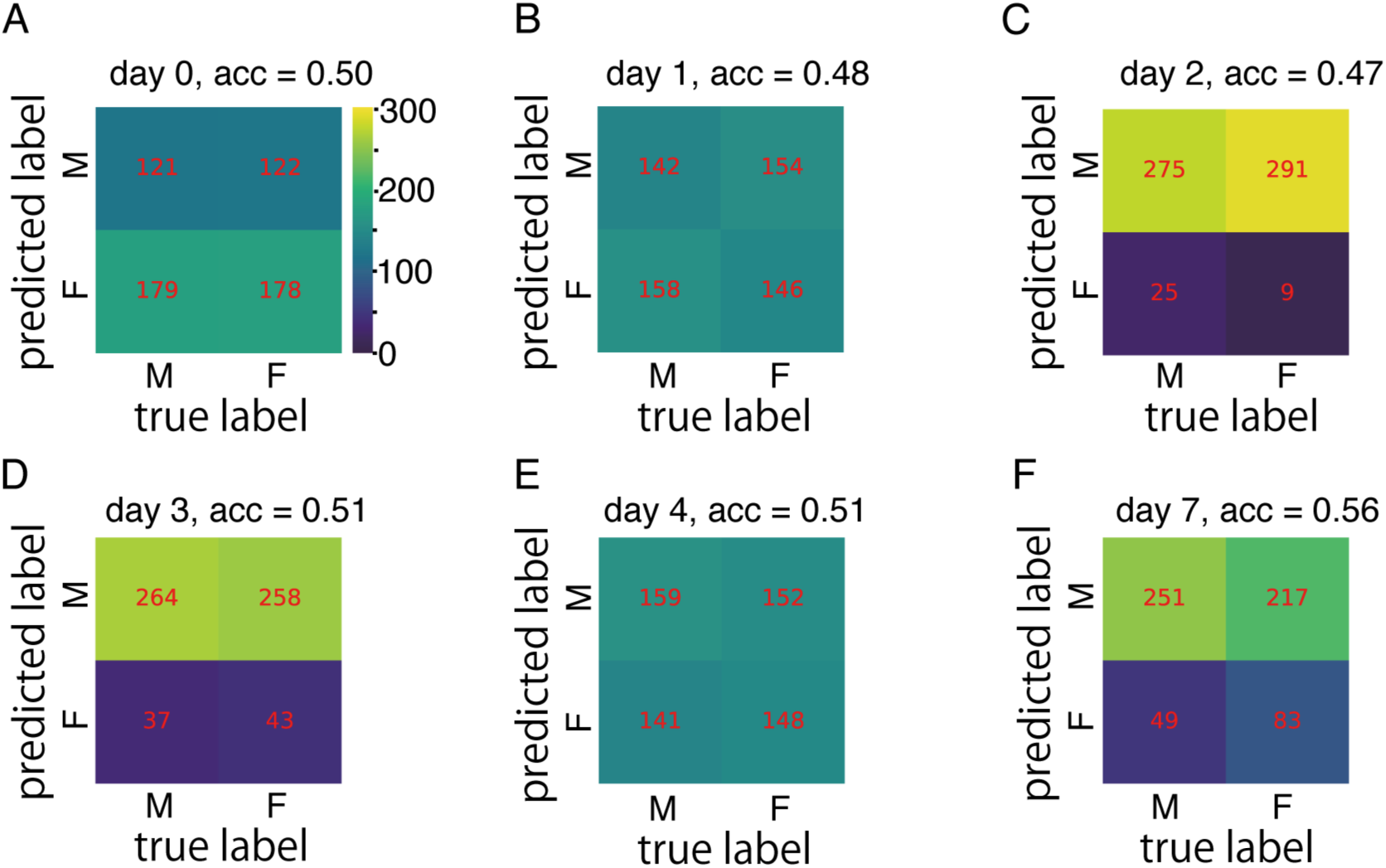
Prediction performance of Tak-1/Tak-2 classifiers on Rit-1/Rit-2 images. (A-F) Confusion matrices showing true sex labels of Rit-1/Rit-2 plants (columns) and predicted sex labels (rows) for day 0 (A), day 1 (B), day 2 (C), day 3 (D), day 4 (E), and day 7 (F). Numbers in red indicate the numbers of Rit-1/Rit-2 images in the corresponding categories. If the prediction is perfectly accurate for both males and females, the matrix should be diagonal. We also computed the prediction accuracy (acc) and Matthew’s correlation constant (MCC) to quantify the model performance.

## Discussion

In this study, we propose a deep learning-based approach to detect and quantify cryptic morphological variability in *Marchantia polymorpha*. We demonstrated the effectiveness of our approach by analyzing the differences between the sexes observed in gemmalings during the first week of development. To distinguish sexual dimorphism from autosomal variations, we established RILs, Rit-1 and Rit-2, from the wild accessions, Tak-1 and Tak-2. To construct a dataset for deep learning, we acquired bright-field images of gemmalings of Rit-1/Rit-2, Tak-1/Tak-1, and male/female of another wild accession, Aus. We observed that deep learning-based ResNet50 successfully classified male and female gemmalings of the wild accessions with >90% test accuracy, while it could not reliably predict the sex of the RILs (Fig. 3). This implies that the RILs have lost the apparent sexual dimorphism present in parental wild accessions, indicating that the morphological differences between Tak-1 and Tak-2 arise from autosomal variations rather than sex chromosomes. Notably, the parental Tak-1 and Tak-2 had identical sequences in >50% of their genomes (Fig. 1), which shows that Tak-1 and Tak-2 had been already inbred at the time of collecting, despite visible morphological differences even within the first week of gemmaling growth. This also highlights the importance of carefully constructed inbred lines; thus, Rit-1 and Rit-2 serve as an appropriate model system for investigating sexual dimorphism. Our approach is particularly important for quantifying the degree of phenotypic loss resulting from genetic changes, because the deep learning-based classifiers eliminate the need for manual feature definition, thus bypassing the “devil’s proof” that manually-defined features may be inadequate for capturing phenotypic changes.

To validate whether the models learned to use features of biological importance, we adopted feature ablation, such as color binarization, and an XAI method, Grad-CAM (Figs. 4 and 5). As a result, we confirmed that after a few days of growth the sex classifications of the trained models depended on the plant body rather than irrelevant features such as scratches on the medium surface. An interesting research direction would be to identify the exact morphological features responsible for this classification. Unfortunately, the Grad-CAM heatmaps do not have sufficient spatial resolution for this task. Higher-resolution visual explanation methods, such as LWRP (Bach et al. 2015) and XRAI (Kapishnikov et al. 2019), are candidates for investigating sex-specific phenotypes in further detail. Another promising research direction is the so-called “Human-in-the-Loop” approach. For example, in the CARTA framework (Kutsuna et al. 2012), a model optimizes a set of features based on human feedback, simultaneously leading to high accuracy and interpretability.

Our approach based on the classifier has the advantage of being applicable to other modalities, such as fluorescence microscopy and infra-red thermography. In particular, recent advancements in fluorescence/luminescence reporters have allowed us to acquire diverse image data for which optimal methods of characterization and quantification have not yet been established. The procedures developed here should facilitate a wide range of applications of deep learning-based methods in plant biology.

## Materials and Methods

### Establishment of the recombinant inbred lines of Marchantia polymorpha

The accessions Tak-1 and Tak-2 (Ishizaki et al., 2008) were used as original female and male strains, respectively, for the generation of recombinant inbred lines. All plants of Tak-2, Tak-1, and their offsprings were cultured on half-strength Gamborg’s B5 (1/2 B5) medium (Gamborg et al., 1968) with 1% agar under continuous light (50 ∼60 µmol photons m^-2^ s^-1^) at 22°C.

To cross male and female plants, gemmae were aseptically incubated on 1/2 B5 medium for more than 2 weeks, then incubated in a container, ECO2box (E1654, DUCHEFA BIOCHEMIE B.V., Netherland), containing 1/2 B5 medium under sexual-organ induction conditions, i.e., continuous white light (50∼60 µmol photons m^-2^ s^-1^) supplemented with far-red light (30 µmol photons m-2 s-1) at 18 °C. After archegoniophores and antheridiophores had been formed, sperm was collected from antheridia as suspension in sterile water and put onto archegoniophores three times a week until sporangia were visibly recognized. Matured sporangia were collected and dried for 1 week at room temperature. Dried sporangia were mixed into sterilized water, then spread onto 1/2 B5 medium plate and grown to the optimal size that is enough for genotyping of sex diagnosis as described previously (Iwasaki et al., 2021).

The spores obtained by the cross between Tak-1 and Tak-2 were designated as the F1 generation. More than ten male and female F1 individuals were selected. Crosses between more than five sibling couples were performed, and one normal-looking sporangium was collected from each cross. These sporangia were separately spread onto 1/2 B5 medium, and one with enough number of spores and a high germination rate was selected as the F2 generation. These procedures were repeated for several generations. A pair of F5 siblings were defined as the male and female recombinant inbred lines Rit-1 and Rit-2.

### Genome sequencing

Rit-1 and Rit-2 genomic DNAs were purified as previously described (Bao et al. 2023). DNA libraries for sequencing were constructed using NEBNext Ultra II FS DNA Library Prep Kit for Illumina and used to obtain paired-end reads using the Illumina NextSeq500 platform.

### Validation of recombinant inbred lines by polymorphism detection

Fastq files available from SRA-run ID DRR120994 were used for the analysis of Tak-2 polymorphisms. The fastq files of Tak-2, Rit-1 and Rit-2 were trimmed by fastp (v0.12.4) (Chen et al. 2018), then mapped against Tak-1 geneme v6.1 (downloaded from MarpolBase [marchantia.info]) by bwa-mem2 (v2.2.1) (Vasimuddin et al. 2019) at default parameters. Output sam files were sorted by samtools (v1.11) (Li et al. 2009) sort, followed by samtools fixmate with -m option. Samtools sort with -n option was used to sort these bam files before samtools markdup to produce dedupped bam files for following variant calling. Due to the lack of a golden standard list of single nucleotide polymorphisms (SNPs) in M. polymorpha unlike human, we usedvariantscalled by GATK (v4.1.3.0) (McKenna et al. 2010) to recalibrate bam files for the second variant call. The first variant call was done by gatk HaplotyoeCaller with -pairHMM LOGLESS_CACHING -ERC GVCF -ploidy 1 options followed by the run of GenotypeGVCF with -ploidy 1 option. Hard filters were applied for each SNP and INDEL to filter out low-quality variants by gatk VariantFilteration and SelectVariants —exclude-filtered. For SNP filtering, QD<2.0, QUAL<30.0, SOR>3.0, FS>60.0, MQ<40.0, MQRankSum < -12.5, and ReadProRankSum < -8.0 criteria were used. QD<2.0, QUAL<30.0, and FS>20.0 filters were used for INDEL filtering. Output vcf files for SNPs and INDELs were fed to gatk BaseRecalibrator to generate a recalibration table with which gatk ApplyBQSR recalibrated bam file for the second variant call. The second variant call was done as described above to gain vcf files for SNPs and INDELs respectively. Detected variants were also filtered out by the threshold described in the first step of variant filtration. To produce circos plot, we used bcftools (v1.9) (Danecek et al. 2021) norm to normalize the called variants and bcftools isec to detect intersections among Tak-2, Rit-1 and Rit-2. A home-made R script (https://github.com/PMB-KU/Rit-dev) was used to count polymorphisms, including SNPs and INDELs, in every 100 kb or 1,000 kb window. The regions that contained more than 100 polymorphisms per 100 kb or 1,000 polymorphisms per 1,000 kb were defined as those with a significant number of polymorphisms in Supplemental Table 1 or Figure 1B, respectively. The circos plot was visualized by the shinyCircos package (Yu, Ouyang, and Yao 2018).

The fasta files of the Rit-1 and Rit-2 genome sequences were created based on SNPs and INDELs using the gatk FastaAlternateReferenceMaker. Filtered reads were mapped against the created genome sequences using bwa-mem2 with default parameters. Coverage was calculated using the bedtools (v2.30.0) (Quinlan and Hall 2010) genomecov, and regions with 0 coverage were masked with N using the bedtools maskfasta. These masked fasta files were registered as PRJDB15748 in International Nucleotide Sequence Database Collaboration.

### Image acquisition

Tak-1/Tak-2, Rit-1/Rit-2, and male and female Aus accessions were cultured from gemmae on 1/2 B5 medium containing 1.0% agar at 22oC under continuous white light. Images were acquired by the digital microscope KH-7700 (HIROX) equipped with the lens MXG-2016Z (day 0∼3: ×60, day 4: ×40, day 7: ×20). The dataset includes 199-200 plants for Aus and Tak-1/Tak-2, and 600-602 for Rit-1/Rit-2, with the same numbers of female and male individuals, except for 99 females and 100 males in Aus 7-day-old gemmalings (see Supplementary Table S2 for details). The timepoints were for days 0, 1, 2, 3, 4, and 7 after planting gemmae.

### Nonlinear PCA of gemmaling images

For data visualization of the gemmaling images, we first pre-processed the images with ImageNet-pretrained ResNet50 (ResNet50_Weights.IMAGENET1K_V2 from the torchvision library) as a feature extractor. Specifically, we extracted image features from the last global average pooling (GAP) layer; hence, the original dataset images (1200 × 1600 × 3) were reduced to 2048-dimensional vectors. Then, for t-SNE clustering, we used a python implementation from the scikit-learn library, “sklearn.manifold.TSNE” (random_state: 1000000, perp: 30, n_iter: 1000).

### Training and validation of deep-learning models

We pre-processed images before feeding them to deep-learning models using the “transform” function in the torchvision library from the PyTorch project (See Supplementary Table S4 for the versions of the libraries used in this study). To construct the training dataset, we began by center-cropping the original images with a crop size of 1200 pixels to remove the excess background regions. Next, we erased the scale bar and augmented the images by flipping and rotating them. Finally, we resized the images to 300 × 300 pixels. The pixel intensities were transformed into the range [0, 1] and normalized using mean [0.485, 0.456, 0.406] and standard deviation [0.229, 0.224, 0.225]. For the validation/test dataset, we used the same pre-processing steps except that we did not apply flipping or rotation.

We used ImageNet-pretrained ResNet50 (ResNet50_Weights.IMAGENET1K_V2 from the torchvision library) as the base model. The penultimate layer was customized for binary classification and only the parameters of the layer were trained. The images were randomly split into training, validation, and test sets at a ratio of 64:16:20. We adopted the Adam optimizer (learning rate: 0.001, batch size: 32, epochs: 500) and used the parameters that achieved the maximum accuracy on the validation set during training.

We adopted the scikit-image library to implement the image modifications in Fig. 3. First, we blurred the images using a Gaussian filter (bandwidth = 10 pixels) and created binary masks using Otsu’s thresholding. Then we applied dilation and hole-filling operations in order to make the masks cover the entire regions of gemmalings. Masks were used to eliminate the background/foreground information in the images. A Gaussian filter (bandwidth = 60 pixels) was applied for severe blurring.

### Visual explanation based on XAI method

We adopted the Grad-CAM method to visualize highly relevant regions for classification of the trained models (Selvaraju et al. 2017). For all test images, we computed the Grad-CAM heatmaps on the last convolutional layer based on a PyTorch implementation (https://github.com/jacobgil/pytorch-grad-cam). Then, to select a representative heatmap for each accession/sex/developmental day, we used the unnormalized logits, i.e., the raw output of the last fully-connected layer in ResNet50. These logits were used to quantify the degree of representativeness of the input images, following recent studies on out-of-distribution detection in deep-learning models (Hendrycks et al. 2019; Vaze et al. 2021).

## Data availability

Image data used in this study have been deposited in the RIKEN SSBD: repository (Systems Science Biological Dynamics repository) with the https://ssbd.riken.jp/repository/290/. Our code used for training of neural networks is available at https://github.com/nyunyu122/Marchantia_sex_classifier. Sequence data are available at ToBeInserted. Lines generated during this study will be shared on reasonable request to Takayuki Kohchi.

## Acknowledgments

We thank John L. Bowman (Monash University) for sharing Australian/Melbourne accessions, Yoshihiro Yoshitake, Yoriko Matsuda, and Chikako Inoue for preparing genomic DNAs, Takafumi Kondo and Yukari Sando for NGS analysis. Computations were partially performed on the NIG supercomputer at ROIS National Institute of Genetics. We also thank the Model Plant Section, Model Organisms Facility, and the NIBB Trans-Scale Biology Center for their technical support.

## Author contribution

Conceptualization: TU, YK. Resources: AK. Investigation: NM, TH. Formal Analysis: YT, ES, SK. Software: YT. Supervision: TK, TU, YK. Writing - original draft: YT, YK, NM. Writing - review and editing: TK, TU, RN. Funding acquisition: TU, TK.

