## Supplementary data for "Harnessing Deep Learning to Analyze Cryptic Morphological Variability of *Marchantia polymorpha*"

**Short title**

deep learning analysis for cryptic morphology

**Corresponding author**

Yohei Kondo

Quantitative Biology Research Group, Exploratory Research Center on Life and Living Systems (ExCELLS), National Institutes of Natural Sciences, 5-1 Higashiyama, Myodaiji-cho, Okazaki, Aichi, 444-8787, Japan

Department of Basic Biology, School of Life Science, SOKENDAI (The Graduate University for Advanced Studies), 5-1 Higashiyama, Myodaiji-cho, Okazaki, Aichi, 444-8787, Japan

**Supplementary Figure S1.** (A) Quantification of overlap between Grad-CAM heatmap and the background by IoU score. The Grad-CAM heatmaps were computed for the predicted sex labels. Each dot represents an image. For each accession and day, the IoU scores for all test images were shown. The color code is the same as that in Fig. 4A. (B) Quantification of overlap between Grad-CAM heatmap and whole image.

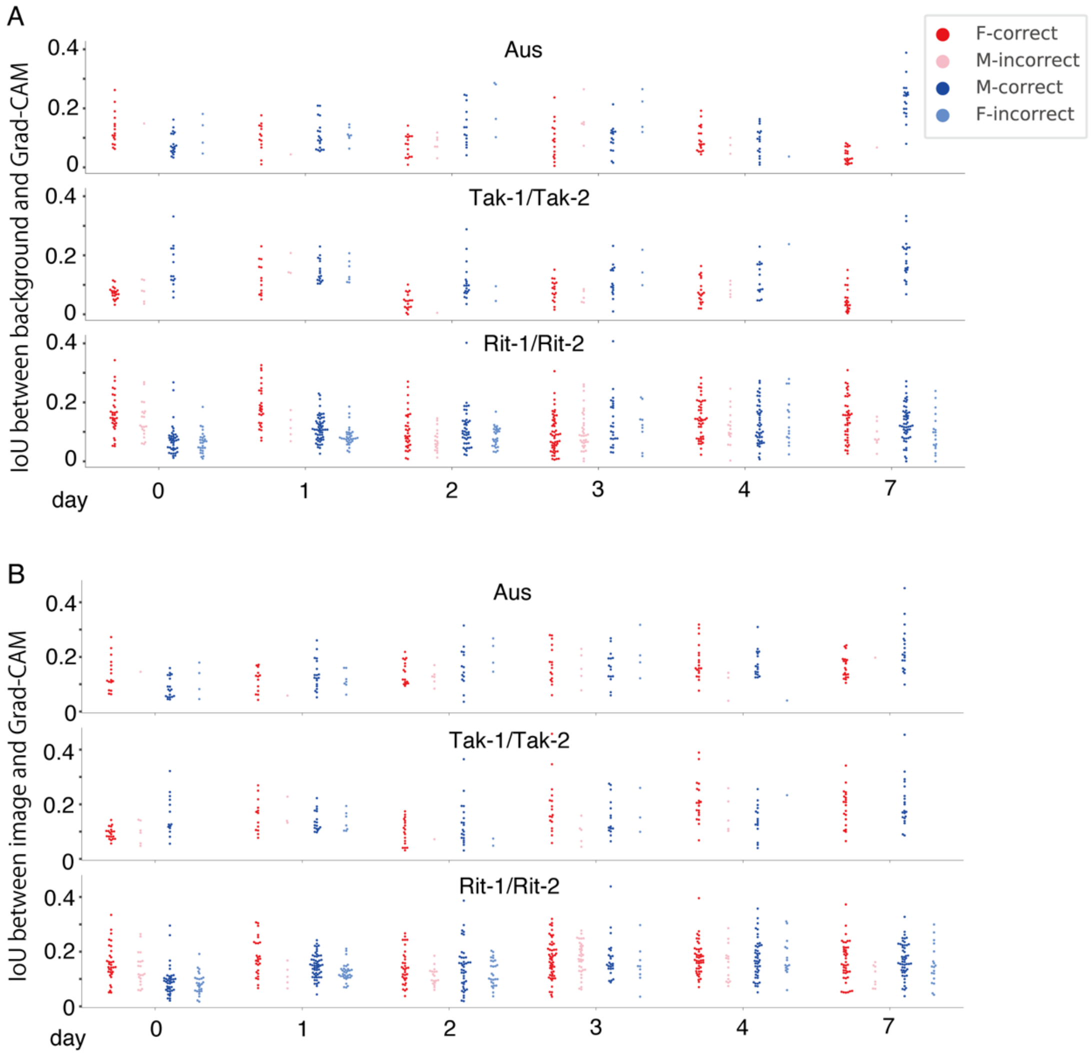

**Supplementary Table S1.** Estimated ratios of autosomal regions derived from Tak-1 or Tak-2.

| | Tak-1-derived regions | | Tak-2-derived regions | | Common regions* | | Whole length |
|---|---|---|---|---|---|---|---|
| | Length (Mb) | Ratio (%) | Length (Mb) | Ratio (%) | Length (Mb) | Ratio (%) | (Mb) |
| Chr 1 | 11.5 | 37.6 | 11.7 | 38.2 | 7.4 | 24.2 | 30.6 |
| Chr 2 | 0.4 | 1.3 | 8.5 | 28.6 | 20.8 | 70.0 | 29.7 |
| Chr 3 | 0.0 | 0.0 | 20.1 | 73.9 | 7.1 | 26.1 | 27.2 |
| Chr 4 | 8.7 | 32.2 | 4.6 | 17.0 | 13.7 | 50.7 | 27.0 |
| Chr 5 | 0.1 | 0.4 | 3.5 | 13.1 | 23.2 | 86.6 | 26.8 |
| Chr 6 | 0.3 | 1.3 | 0.1 | 0.4 | 23.5 | 98.3 | 23.9 |
| Chr 7 | 4.6 | 20.9 | 3.8 | 17.3 | 13.6 | 61.8 | 22.0 |
| Chr 8 | 0.1 | 0.5 | 11.7 | 54.7 | 9.6 | 44.9 | 21.4 |
| Total | 25.7 | 12.3 | 64.0 | 30.7 | 118.9 | 57.0 | 208.6 |

The lengths were estimated from the number of 100-kb windows in each category.

Asterisk indicates regions shared between Tak-1 and Tak-2.

**Supplementary Table S2. Number of images.**

See the file “Supplementary Table S2.xlxs”.

**Supplementary Table S3. Effect size of between the sexes, calculated as Hedge’s g*.**

| strain | day | unbiased Cohen’s d |
|---|---|---|
| Tak-1/Tak-2 | 0 | 1.3116 |
| Tak-1/Tak-2 | 1 | 1.3929 |
| Tak-1/Tak-2 | 2 | 1.7103 |
| Tak-1/Tak-2 | 3 | 1.5961 |
| Tak-1/Tak-2 | 4 | 1.4969 |
| Tak-1/Tak-2 | 7 | 2.0090 |
| Aus | 0 | 0.4156 |
| Aus | 1 | 0.2109 |
| Aus | 2 | 0.3093 |
| Aus | 3 | 0.0209 |
| Aus | 4 | 0.3348 |
| Aus | 7 | 1.6032 |
| Rit-1/Rit-2 | 0 | 0.2208 |
| Rit-1/Rit-2 | 1 | 0.1759 |
| Rit-1/Rit-2 | 2 | 0.0692 |
| Rit-1/Rit-2 | 3 | 0.0231 |
| Rit-1/Rit-2 | 4 | 0.1197 |
| Rit-1/Rit-2 | 7 | 0.1205 |

**Supplementary Table S4. Versions of the libraries used in this study.**

| |
|---|
| Python: 3.6 (in anaconda)<br>PyTorch: 1.7.1<br>Torchvision: 0.8.2 |
| Ubuntu: 16.04<br>Machine: DeepLearning BOX II, 64 GB RAM, GPU=4, GeForce RTX 2080 Ti<br>CUDA driver: 430.34<br>CUDA: 10.1 |
| numpy: 1.18.5<br>scikit-image: 0.16.2<br>scikit-learn: 0.23.1 |

Supplementary Table S2. Number of images.

| accession | day | sex | hue | number of images |
|---|---|---|---|---|
| Aus | 0 | M | Aus_0d_M | 100 |
| Aus | 0 | F | Aus_0d_F | 100 |
| Aus | 1 | M | Aus_1d_M | 100 |
| Aus | 1 | F | Aus_1d_F | 100 |
| Aus | 2 | M | Aus_2d_M | 100 |
| Aus | 2 | F | Aus_2d_F | 100 |
| Aus | 3 | M | Aus_3d_M | 100 |
| Aus | 3 | F | Aus_3d_F | 100 |
| Aus | 4 | M | Aus_4d_M | 100 |
| Aus | 4 | F | Aus_4d_F | 100 |
| Aus | 7 | M | Aus_7d_M | 100 |
| Aus | 7 | F | Aus_7d_F | 99 |
| Tak-1 | 0 | M | Tak-1_0d_M | 100 |
| Tak-2 | 0 | F | Tak-2_0d_F | 100 |
| Tak-1 | 1 | M | Tak-1_1d_M | 100 |
| Tak-2 | 1 | F | Tak-2_1d_F | 100 |
| Tak-1 | 2 | M | Tak-1_2d_M | 100 |
| Tak-2 | 2 | F | Tak-2_2d_F | 100 |
| Tak-1 | 3 | M | Tak-1_3d_M | 100 |
| Tak-2 | 3 | F | Tak-2_3d_F | 100 |
| Tak-1 | 4 | M | Tak-1_4d_M | 100 |
| Tak-2 | 4 | F | Tak-2_4d_F | 100 |
| Tak-1 | 7 | M | Tak-1_7d_M | 100 |
| Tak-2 | 7 | F | Tak-2_7d_F | 100 |
| Rit-1 | 0 | M | Rit-1_0d_M | 300 |
| Rit-2 | 0 | F | Rit-2_0d_F | 300 |
| Rit-1 | 1 | M | Rit-1_1d_M | 300 |
| Rit-2 | 1 | F | Rit-2_1d_F | 300 |
| Rit-1 | 2 | M | Rit-1_2d_M | 300 |
| Rit-2 | 2 | F | Rit-2_2d_F | 300 |
| Rit-1 | 3 | M | Rit-1_3d_M | 301 |
| Rit-2 | 3 | F | Rit-2_3d_F | 301 |
| Rit-1 | 4 | M | Rit-1_4d_M | 300 |
| Rit-2 | 4 | F | Rit-2_4d_F | 300 |
| Rit-1 | 7 | M | Rit-1_7d_M | 300 |
| Rit-2 | 7 | F | Rit-2_7d_F | 300 |
